# Cardiac-cerebrovascular crosstalk: Cardiac rhythms reveal maladaptive cerebral blood flow velocity and constrained ventilatory status

**DOI:** 10.1101/2025.10.21.683641

**Authors:** Diego Candia-Rivera, Pierre Pouget, Mario Chavez

## Abstract

In brain-heart interactions, several pathways have been proposed to mediate feedback loops between systems. Among these, cerebrovascular dynamics operate at their interface. However, how cardiovascular control, ventilation mechanisms, and cerebral autoregulation interact is not well characterized, especially in ageing and post-stroke conditions, where perfusion can be compromised. In a cohort of 57 elderly participants, including 30 stroke survivors, we investigated the relationship between cardiac sympathetic activity and both, cerebral blood flow regulation and ventilatory status. Sympathetic reflexes, assessed via cardiac sympathetic index (CSI) during sit-to-stand transitions, were preserved across all participants, with marginal group differences between stroke and non-stroke populations. However, among individuals with constrained ventilation, indexed by reduced end-tidal CO₂ at baseline, we identified a more elevated CSI following postural change, scaling with the degree of CO₂ dysregulation. Furthermore, transcranial Doppler measurements revealed exaggerated changes in mean flow velocity (MFV) within the right middle cerebral artery in most participants. These MFV shifts significantly correlated with the magnitude of cardiac sympathetic change under orthostatic stress, suggesting that CSI can capture maladaptive cerebrovascular responses. Together, these findings highlight a distinct cardiac-cerebrovascular crosstalk in elderly individuals, revealing patterns consistent with compensatory or maladaptive sympathetic overactivation under conditions of impaired cerebrovascular control.

## Introduction

Understanding the interplay between cardiac and cerebrovascular regulation is essential for grasping how the brain and bodily systems adapt to physiological stressors. While traditionally considered as distinct systems, recent research into brain-heart interactions increasingly supports a view of bidirectional communication between neural and cardiovascular systems [1], where cerebrovascular dynamics appear at the interface [2,3], although the extent of direct neural control over cerebral blood flow remains debated [4,5]. This cardiac-cerebrovascular crosstalk becomes particularly relevant in dynamic conditions such as postural changes, where both autonomic cardiovascular responses and cerebrovascular adjustments must occur in concert [6]. Despite this emerging integrative view [7], the specific ways in which cardiac autonomic output relates or interacts with cerebrovascular regulatory mechanisms remain insufficiently characterized [8].

Cerebral autoregulation is the brain’s intrinsic mechanism to maintain stable cerebral blood flow in the face of systemic blood pressure fluctuations. This complex control system involves myogenic, metabolic, and neurogenic inputs and is essential for maintaining optimal perfusion during physiological challenges [6,9]. Superimposed on these local mechanisms, especially under stress conditions, autonomic pathways can also intervene through sympathetic efferent projections to regulate vasoconstriction, blood pressure and heart rate. Although cerebral autoregulation is primarily mediated by small resistance vessels, beat-to-beat changes in middle cerebral artery (MCA) mean flow velocity provide a surrogate of downstream autoregulatory activity [10]. The influence of sympathetic nervous activity on cerebral blood flow, however, remains debated [4]. In humans, sympathetic effects may contribute to dynamic autoregulation in a frequency-dependent manner [11–13]. Anatomical pathways linking autonomic nuclei to both cardiac efferent and large cerebral vessels, including sympathetic projections relayed via the superior cervical ganglion to the MCA, provide a substrate for partially coordinated regulation of heart function and cerebral blood flow [14]. Complementary baroreceptor and chemoreceptor afferents projecting to the nucleus tractus solitarius further integrate cardiovascular and cerebrovascular control, shaping autonomic outflow to both systems.

Arterial CO_2_ is a potent regulator of cerebrovascular tone and a primary driver of both static and dynamic components of cerebral autoregulation [15–17]. Because direct measurement of arterial CO_2_ is invasive, end-tidal CO_2_ (EtCO_2_) is commonly used as an accessible surrogate [18]. Reductions in EtCO₂ may reflect impaired ventilatory drive, increased physiological dead space, or inefficient gas exchange, all of which have been associated with heightened cerebrovascular vulnerability in older adults and in disease populations.

In parallel, autonomic dynamics during orthostatic stress have been explored as markers of altered cerebrovascular regulation [19–22]. Prior work has often examined these autonomic signals in isolation, without considering their interaction with cerebrovascular mechanisms. Yet cardiac activity is embedded within central neural circuits: converging evidence demonstrates mutually influential brain-heart loops, in which central autonomic networks modulate cardiac rhythms and respond to afferent cardiovascular feedback [23–25].

Here, we refer to cardiac–cerebrovascular crosstalk as the coordinated interaction between cardiac autonomic rhythms and cerebrovascular regulatory processes, including autoregulatory adjustments, vascular reactivity, and potentially neurovascular coupling. For clarity, the present study focuses on three interacting physiological domains: (i) cardiac autonomic activity (indexed by CSI), (ii) cerebrovascular regulation (indexed by MCA blood flow velocity), and (iii) ventilatory status (indexed by EtCO_2_), and examines their pairwise and joint relationships under orthostatic stress. Under this framework, maladaptive cerebrovascular control may be detectable through altered cardiac autonomic signatures, particularly during dynamic challenges such as orthostatic transitions.

To capture these rapid autonomic adjustments, we recently developed a method for time-resolved estimation of heart rate dynamics, from which the cardiac sympathetic index (CSI) is derived [26]. Sympathetic responses ensure adequate cerebral perfusion during postural shifts by modulating heart rate, vascular tone, and cardiac output [27]. Although an intact autonomic state is acknowledged as necessary for optimal cerebral autoregulation [8], commonly used autonomic markers such as heart rate variability (HRV) have not been systematically applied to characterize cerebral autoregulation, in part because the physiological origins of HRV spectral components remain debated [28]. Evidence supporting autonomic relationships with cerebral autoregulation includes reports of patient groups with impaired autoregulation who exhibit altered HRV spectral patterns [29,30]. In addition, reduced baroreflex sensitivity, typically accompanied by relatively greater sympathetic drive, has been associated with stronger autoregulatory control [29,31]. However, given the uncertainties of HRV spectral interpretation, we sought to examine cardiac–cerebrovascular relationships using CSI, which provides a more direct and validated estimate of sympathetic activity [26].

Despite the parallel importance of cerebral autoregulation and cardiac autonomic control, relatively little work has integrated these domains in the context of ageing and stroke [29,30]. Stroke may disrupt central autonomic pathways, damage cerebrovascular networks, and create asymmetries in neurovascular control, especially when affecting key cortical or subcortical regions involved in motor and autonomic integration [32]. It remains unclear whether impaired cerebrovascular regulation in stroke is associated with changes in sympathetic outflow, or whether the systems become uncoupled altogether, leading to maladaptive physiological patterns.

We therefore hypothesized that cardiac–cerebrovascular crosstalk would provide an integrated signal of impaired cerebral blood flow regulation, particularly in individuals exhibiting constrained ventilatory status, indexed by reduced EtCO_2_. Specifically, we aimed to determine whether cardiac sympathetic activity is differentially associated with cerebrovascular regulation depending on ventilatory status, and whether such relationships reflect altered physiological coordination under orthostatic stress. We expected distinct cardiac sympathetic reflexes in individuals with constrained CO_2_ profiles, as well as altered cerebral blood flow velocity responses to postural changes.

To test these hypotheses, we investigated how cardiac sympathetic activity, cerebral blood flow velocity, and ventilatory status interrelate in a cohort of 57 elderly individuals, including 30 stroke survivors. Using a postural change test (Figure 1), we quantified sympathetic responsiveness through CSI and examined its relationship to both resting EtCO_2_ levels and cerebral blood flow velocity changes, assessed via transcranial Doppler ultrasound of the middle cerebral arteries. We further explored whether these relationships differed between stroke and non-stroke participants. These analyses were designed to assess the value of monitoring cardiac–cerebrovascular crosstalk as a marker of physiological vulnerability in ageing and stroke.

**Figure 1.**
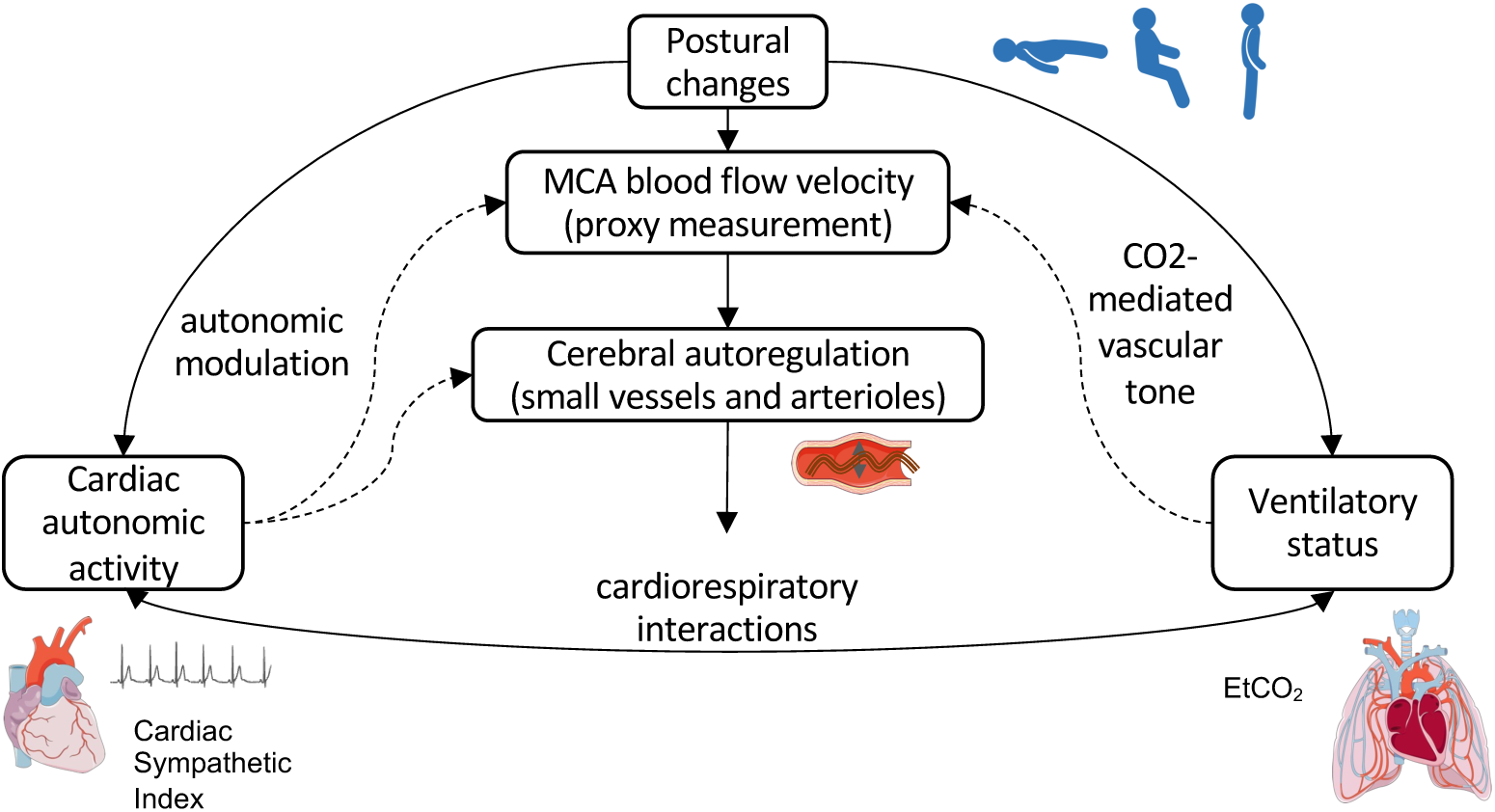
Conceptual framework of cardiac–cerebrovascular interactions under postural changes. The study examines interactions between three physiological domains: cardiac autonomic activity (indexed by the cardiac sympathetic index, CSI), cerebrovascular regulation (indexed by middle cerebral artery, MCA, blood flow velocity), and ventilatory status (indexed by end-tidal CO_2_, EtCO_2_). Cerebral autoregulation primarily occurs at the level of small resistance vessels and arterioles, which adjust vascular tone to maintain stable cerebral perfusion. Transcranial Doppler measurements of MCA blood flow velocity provide a non-invasive proxy of these downstream autoregulatory processes but do not directly measure the site of regulation. Arterial CO_2_ is a major modulator of cerebrovascular tone and reactivity, influencing autoregulatory capacity, whereas autonomic influences on cerebral blood flow are indirect and remain debated. Orthostatic stress (supine-to-sit and sit-to-stand transitions) engages these systems simultaneously, enabling the assessment of their coordinated responses. Dashed arrows represent hypothesized associations and physiological interactions rather than direct causal pathways.

## Materials and methods

### Participants

The data used in this study belongs to the Physionet dataset “Cerebral Vasoregulation in Elderly with Stroke” [33,34]. Participants were recruited in Boston, USA, from community advertisement. The inclusion criteria aimed at recruiting participants aged 60-80 years, both stroke and non-stroke (control) groups. In this study, a total of 57 participants were analysed (61 participants were available, from which 4 were excluded because their corrupted ECG during the analysed protocol due to artifacts), from which 30 correspond to stroke survivors.

The control group inclusion required no clinical history of stroke and no focal deficit on neurological exam. The stroke group inclusion consisted in patients with at least 6 months from the stroke and clinically stable condition (defined by neurological scales Modified Rankin Scale, MRS<4; and moderate National Institute of Health Stroke Scale, NIHSS <15, see Supplementary Table 3 and Supplementary Figure 2). Inclusion also requested patients with first ever hemispheric ischemic stroke with documented neurological deficit persisting >24 hours, confirmed by radiological findings (CT and MRI), and infarct affecting less than one third of the middle cerebral artery territory. In the present study, patients presented different types of aetiologies, being atherosclerosis the most common (14). The other aetiologies included cardioembolism (3), atherothrombosis (3), other embolic (2) and unknown aetiology (8). Five patients had subcortical lesions. Five patients had lesions in known nodes of the central autonomic network (insula, thalamus, basal ganglia), although it is not known in all patients whether lesions in the frontal lobe reached relevant structures, such as the insula or the anterior cingulate cortex (part of the anterior artery territory). See Supplementary Table 1 for detailed aetiologies, affected hemispheres and regions.

Exclusion criteria for the control group included individuals with carotid stenosis greater than 50%, based on clinical history, ultrasound, or magnetic resonance angiography (MRA). For the stroke group, subjects were excluded if they had carotid stenosis greater than 50% that was contralateral to the stroke or not related to the infarct, as well as those with evidence of intracranial or subarachnoid haemorrhage on MRI or CT. Additional exclusions for the stroke group included the presence of any acute or unstable medical condition; vertebrobasilar or carotid artery disease identified by MRA during initial evaluation; a history of dementia or cognitive impairment that limited compliance with the study protocol; diabetes mellitus; hemodynamically significant valvular disease; clinically significant arrhythmias such as atrial fibrillation; severe hypertension, defined as systolic blood pressure > 200 mmHg and/or diastolic blood pressure >110 mmHg, or the use of three or more antihypertensive medications (see Supplementary Table 2 for detailed medication information). Participants were also excluded if they had current alcohol or recreational drug abuse, morbid obesity defined as a body mass index >35, uncontrolled hypertension during taper, hypertension tapering was not possible, no insonation window, or if it was not possible to obtain permission for participation from their primary care physician.

### Protocol

The protocol was performed at the Clinical Research Center from the Beth Israel Deaconess Medical Center, Boston, USA. All participants signed an informed consent and completed symptom checklist and medical history questionnaires. ECG and vital signs, height and weight were measured by a trained nurse.

Antihypertensive medications, if any, were tapered during the 5 days prior to the study using home blood pressure monitoring (from the studied cohort, hypertensive medications were tapered in 12 controls and 22 stroke patients, see Supplementary Table 2). Active standing is a standard method for the diagnosing orthostatic hypotension. In elderly people, the normal response to standing up is a transient systolic blood pressure decline with recovery within the first minute [35], which is compensated primarily by increased peripheral resistance. Orthostatic hypotension (hereafter referred as dysautonomia) will be diagnosed using standard criteria accepted by The Consensus Committee of the American Autonomic Society and American Academy of Neurology, as a systolic blood pressure decline of ≥20 mm Hg or diastolic blood pressure decline of ≥10 mm Hg within 3 minutes of standing [36].

The protocol encompassed several tasks. In this study, we focused in three conditions: supine position (hereafter referred as baseline), sitting and standing. The experimental design of the original study, from which this dataset was derived [33,34], defined the supine position as the baseline, which included an stabilization of blood pressure and heart rate. When subjects transitioned to the sitting condition, they sat on the chair with their legs elevated at 90 degrees in front of them on a stool to reduce venous pooling. The moment when both feet touched the ground was recorded to mark the onset of standing. The physiological measurement of the standing condition started after 1-3 minutes from the onset. The subjects were asked to breathe according to a metronome at their normal breathing rate. Physiological measurements were performed for approximately 5 minutes in each condition.

### Data acquisition and processing

This study constitutes a retrospective analysis of a previously published open-access dataset. For transparency, a brief overview of the data acquisition and preprocessing procedures is presented below. Detailed methodological information, including the original study design and experimental protocol, is available in the primary publication [33,34].

Heartbeats were recorded using a 3-lead electrocardiogram. Beat-to-beat arterial pressure was acquired from a cuff placed on the finger on the unaffected side in the stroke group or on the nondominant arm in the control group using a Portapress-2 device (Finapres Medical Systems, Enschede, The Netherlands). Beat-to-beat blood pressure measurements were corroborated by standard Dinamap measurements of arterial pressure on the upper arm (Johnson and Johnson, New Brunswick, NJ, USA). End-tidal CO2 values were recorded using an infrared end-tidal volume gas monitor (Capnomac Ultima, Datex Ohmeda, Chicago, IL, USA) attached to a face mask. A transcranial Doppler ultrasonography system (MultiDop X4, DWL Neuroscan, Sterling, VA, USA) was be used to monitor blood flow velocity in both middle cerebral arteries (see Supplementary Figure 3 for sample of measurements), and in some cases, the radial arteries. The middle cerebral arteries were insonated from the temporal windows, by placing the 2 MHz probe in the temporal area above the zygomatic arch. Each probe was positioned to record the maximal blood flow velocity and fixed at the desired angle using a three-dimensional positioning system attached to the light-metal probe holder. Special attention is always given to stabilize the probes, since their steady position is crucial for continuous blood flow velocity recordings. Fourier transform of the Doppler shift, a difference between the frequency of the emitted signal and its echo (frequency of reflected signal), was used to calculate blood flow velocity. Mean blood flow velocity was detected from the envelope of the arterial flow waveforms. Signals were recorded at either 500 or 1000 Hz. Analog signals were recorded using Labview NIDAQ (National Instruments Data Acquisition System 64 Channel/100 Ks/s, Labview 6i, Austin, TX, USA).

### Cardiac sympathetic activity estimation

ECG time series were bandpass filtered between 0.5 and 45 Hz using a fourth-order Butterworth filter. R-peaks were automatically detected using the Pan-Tompkins algorithm [37]. To identify potential misdetections, an automated procedure flagged sudden changes in inter-beat intervals, which were then reviewed through visual inspection. All detected peaks were subsequently examined manually against the raw ECG signal and the inter-beat interval (IBI) histogram. Manual corrections were applied when necessary, although these were minimal and limited to a few instances.

Cardiac sympathetic activity was assessed using a method based on the time-varying geometry of the IBIs in the Poincaré plot [26]. This approach integrates dynamic changes in heart rate and heart rate variability to derive the Cardiac Sympathetic Index (CSI). The time-resolved computation of CSI was performed using a sliding time window to quantify the fluctuating properties of the successive changes IBIs, depicted as an ellipsoid in the Poincaré plot. The two geometrical features extracted are the long ratio of the ellipsoid (*SD_2_*) and its distance to the origin (*R*). *SD_2_* represents the slow fluctuations in HRV (typically falling within the low frequency range, <0.15 Hz, without the need of a frequency band definition). *R* represents the very slow changes in cardiac cycle duration (CCD), without the low and high-frequency variability components. The two metrics were combined as the Cardiac Sympathetic Index (CSI):

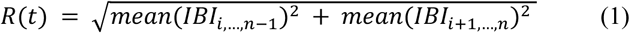

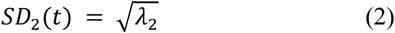

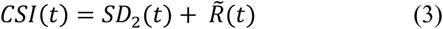

where 𝜆_2_is the larger eigenvalue of the covariance matrix of 𝐼𝐵𝐼_1,…,N-1_ and 𝐼𝐵𝐼_2,…,N_, and 𝑁 is the length of the 𝐼𝐵𝐼 array in the current time window. Before CSI computation, 𝑅(𝑡) and 𝑆𝐷_2_(𝑡) means were re-centred, based on their computation through the whole recording, and R̃ is the flipped 𝑅 with respect the mean. The sliding time window used had a length of 15 s, as per previous simulations in humans [38,39], and a step defined by each heartbeat.

This marker has been validated under standard autonomic challenges [26,40,41], confirming their reliability for capturing time-resolved cardiac autonomic dynamics. For a comprehensive description of the cardiac autonomic estimation procedure, refer to the original method article [26].

### Exposure and outcome measures

The primary exposure variable was the cardiac sympathetic activity, quantified using the CSI, across different physiological and participants’ groups. The primary outcome was the CSI response to an orthostatic challenge (sit-to-stand task), assessed by comparing CSI values during the sitting and standing phases. Group comparisons were performed between controls and stroke patients to evaluate whether the CSI response differed by clinical status. Additional comparisons assessed the influence of dysautonomia, stroke hemisphere, and baseline physiological states.

Associations between CSI and other physiological parameters were examined to identify potential modulators of the sympathetic response. First, we analysed the relationship between CSI and heart rate and blood pressure, as a control, to demonstrate that CSI cannot be trivially equated with standard measures. Specifically, we compared CSI with: (1) baseline blood pressure and heart rate; (2) changes in blood pressure and heart rate during the sit-to-stand task. We then tested CSI in relation to: (1) baseline end-tidal CO₂ (EtCO₂); and (2) percentage changes in cerebral blood flow velocity from baseline to both sitting and standing phases. We focus in the right middle central artery because of its known dominance in autonomic control [42] and postural balance [43–45]. We also focus on the percentual changes in blood flow velocity (baseline with respect to the postural change), to reduce potential confounds associated with individual anatomical differences. Each of these variables was tested for their correlation with CSI within and across groups. These analyses aimed to determine the extent to which cardiac sympathetic modulation is influenced by cardiovascular, respiratory, and cerebrovascular dynamics.

Due to incomplete data availability, as some participants lacked specific physiological measurements, the sample included in each statistical analysis varied slightly across comparisons. Accordingly, participants were stratified according to the availability of the relevant variables for each analysis, and the corresponding sample sizes are detailed in Table 1.

**Table 1.** Participants inclusion in each analysis.

| Condition/measure | Groups' count* |
| --- | --- |
| Dysautonomia | Control=9, stroke=14 |
| No dysautonomia | Control=22, stroke=16 |
| Blood pressure/heart rate | Control=27, stroke=30 |
| CO2 constrained | Control=7, left stroke=5, right stroke=4, stroke no hemisphere information = 1 |
| CO2 unconstrained | Control=20, left stroke=7, right stroke=10, stroke no hemisphere information = 3 |
| Blood flow velocity (right MCA) | Control=9, left stroke=8, right stroke=8, stroke no hemisphere information = 1 |
| Blood flow velocity (left MCA) | Control=18, left stroke=8, right stroke=8, stroke no hemisphere information = 2 |

\* Additional exclusion of patients because of corrupted or missing information on the specific measurement

Further stratification was performed, based on the range of the measurements of EtCO_2_ and blood flow velocity. CO_2_ constrained patients were identified as EtCO_2_ <35 mmHg, as these values are typically associated with overall higher health risk [46–48]. In addition, we performed a data-driven stratification of the cohort by splitting patients into “low” and “high” groups and performed separate linear regressions of CSI versus EtCO₂ within each group. The sum of squared errors (SSE) across both regressions was computed, and the cutoff minimizing the total SSE was identified as the optimal threshold. Typical change in blood flow velocity in healthy individuals is within 10-15% [49,50]. Therefore, we focused on the “abnormal” range 10-100%, to test whether the degree of change in blood flow velocity relates to CSI.

### Statistical analysis

Continuous variables were expressed as medians and median absolute deviations. Paired and unpaired comparisons were performed using Wilcoxon-Mann-Whitney tests. Spearman correlations were used to test relationships between physiological variables. Statistical significance was set to α=0.05.

Marginal differences that did not reach the significance threshold were further evaluated using a bootstrapping approach to estimate the effect size and determine the minimum cohort size at which the observed effect would likely become statistically significant [51].

To obtain a robust, distribution-free effect size, we computed the rank-biserial correlation, which is the canonical effect size for rank-based group comparisons. We estimated its sampling distribution using 10,000 bootstrap iterations. In each iteration, group values were resampled with replacement, the rank-sum statistic was recomputed, and the corresponding rank-biserial effect size was derived (Cliff’s delta, with large effect size considered as δ>0.474 [52]). The resulting bootstrap distribution was used to obtain the mean effect size and the 95% confidence interval.

To estimate the cohort size at which the observed effect would be expected to reach statistical significance (based on Cliff’s delta and estimated mean p-value), we performed a bootstrap subsampling analysis. For a range of hypothetical sample sizes, we randomly drew N samples (with replacement) from each group and recomputed the rank-sum p-value. This procedure was repeated 10,000 times per sample size, yielding an empirical estimate of effect size and p-value, as a function of N. The minimum N at which δ>0.474 or mean p<0.05 were taken as the estimated required sample size for both groups.

### Code and data availability statement

All physiological data used in this study are publicly available in Physionet [33,34]. Codes implementing the methods of this study are available at https://github.com/diegocandiar/robust_hrv.

## Results

We characterized cardiac sympathetic responsiveness during the sit-to-stand task by analysing CSI dynamics in older adults and stroke patients. We first quantified the magnitude and consistency of the CSI across groups, then examined its associations with blood pressure, heart rate, CO_2_ regulation, and cerebral blood flow modulation. These analyses provide an overview of autonomic integrity and its physiological modulators in vulnerable populations.

### Cardiac sympathetic reflex to orthostatic challenge

Figure 2A illustrates the changes in the CSI during a sit-to-stand test in both controls and stroke patients. In both groups, CSI significantly increased during the standing phase compared to the sitting baseline (Figure 2B; Wilcoxon paired test, controls p<0.0001, Z=4.5407, stroke p<0.0001, Z=4.7832). This confirms that stroke patients have intact cardiac sympathetic reflex to orthostatic stress, with no apparent delays with respect to controls. The magnitude of this increase (CSI stand-CSI sit) did not differ significantly between groups (Wilcoxon unpaired test, p=0.9173, Z=0.1039). This trend was also observed in individuals exhibiting orthostatic dysautonomia, indicating that the CSI increase is preserved even in those with impaired blood pressure regulation (Figure 2C; Wilcoxon paired test, controls p=0.0077, Z=2.6656, stroke p=0.001, Z=3.2974). Marginal differences were observed in the magnitude of the sympathetic reflex, where controls had a group-wise larger effect, although not reaching statistical significance (Wilcoxon unpaired test, control vs stroke p = 0.0725, Z=1.7962). This difference was further evaluated using a bootstrapping approach, which yielded an effect size of δ = 0.459 (95% CI [0.016, 0.841]). Based on this, we estimated that a minimum sample size of N = 11 per group would be required to achieve a large effect size (δ > 0.474), and N = 13 per group to reach statistical significance at α<0.05.

**Figure 2.**
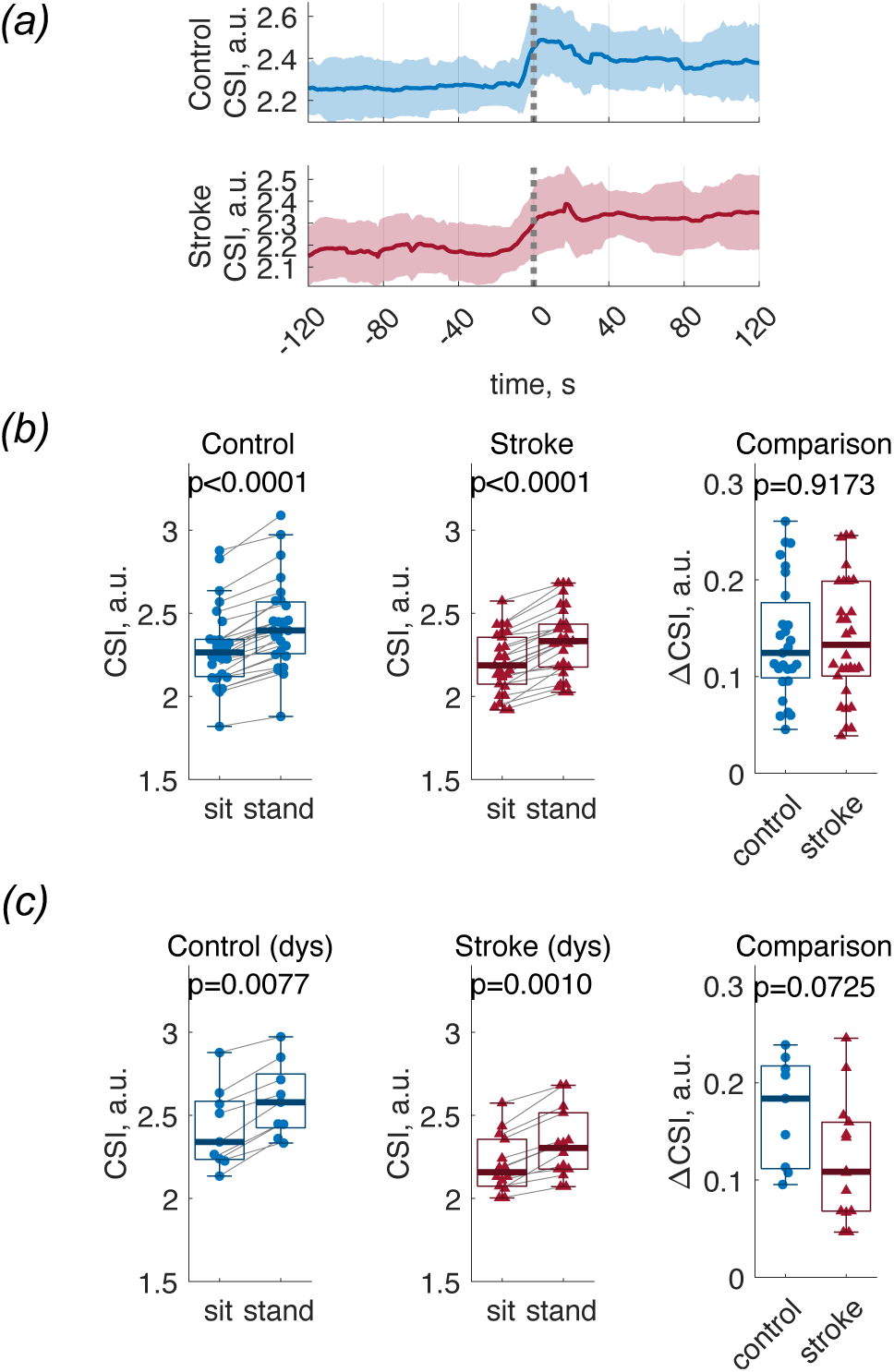
Cardiac sympathetic response to orthostatic stress. (A) Group median CSI continuous values during the sit-to-stand test in controls (n=27) and stroke patients (n=30). (B) Both groups show a significant increase in CSI from sit to stand, with no significant difference in the CSI change (ΔCSI) between groups. (C) The pattern of increased CSI during standing is maintained in individuals with dysautonomia (dys) in both controls (n=9) and stroke patients (n=14).

We then assessed whether the cardiac sympathetic reflex related to other cardiovascular variables. At rest, CSI values showed no significant association with baseline blood pressure (Figure 3A; Spearman Correlation, stroke p=0.3889, r=-0.1626, control p=0.0701, r=-0.3547) or with changes in blood pressure during the sit-to-stand manoeuvre (Figure 3B; Spearman Correlation, stroke p=0.4379, r=-0.1522, control p=0.5607, r=0.1275). A negative correlation was observed between CSI and baseline heart rate in the control group, but not in the stroke group (Figure 3C; Spearman Correlation, stroke p=0.1889, r=-0.2463, control p=0.0099, r=-0.4921). Moreover, CSI changes during the sit-to-stand test were significantly correlated with heart rate changes in both groups (Figure 3D; Spearman Correlation, stroke p=0.0003, r=0.6426, control p<0.0001, r=0.7747), suggesting that although CSI is influenced by heart rate modulation, it also reflects additional autonomic components.

**Figure 3.**
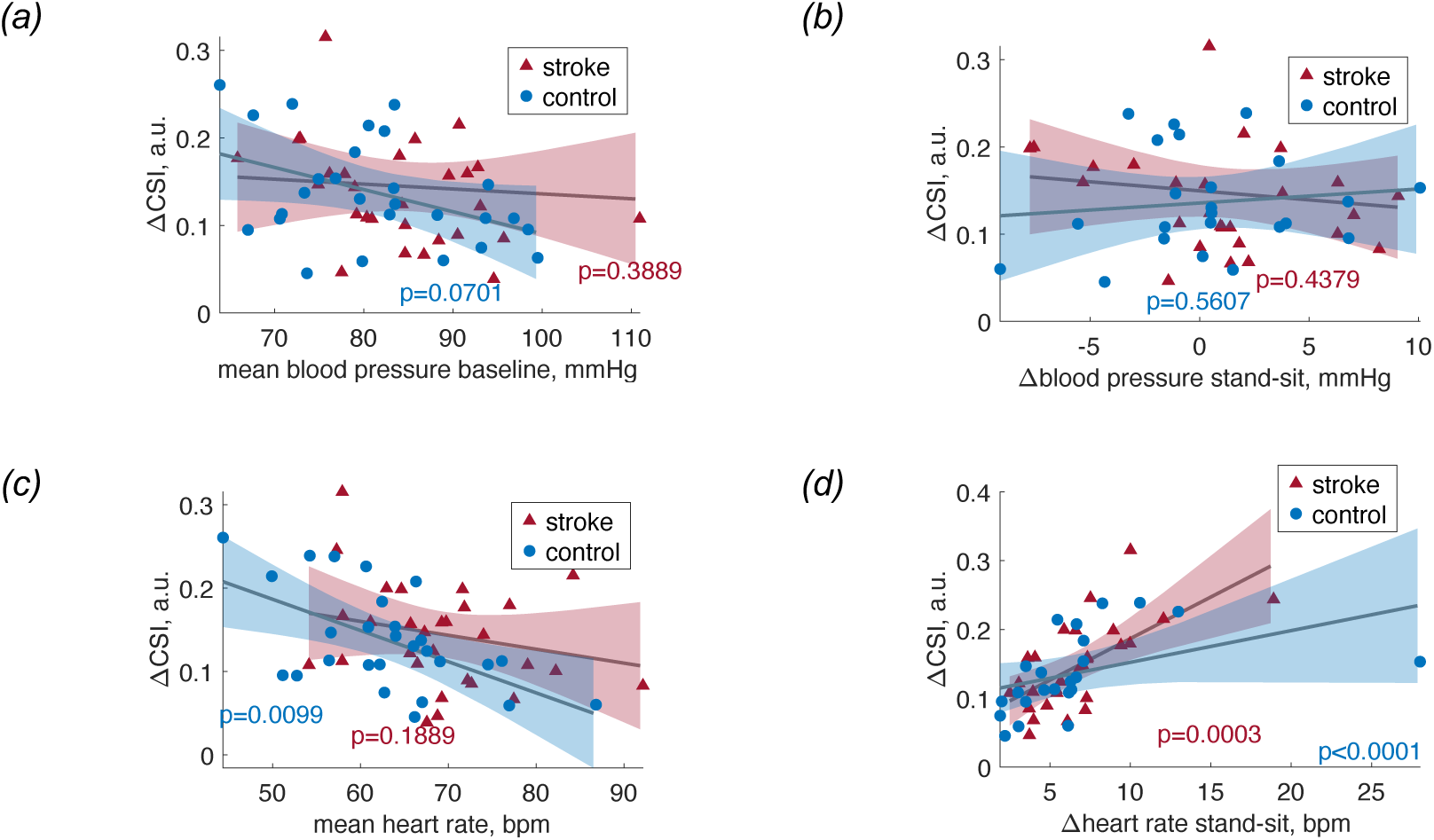
Relationship between cardiac sympathetic reflex and cardiovascular parameters in controls (n=27) and stroke patients (n=30). (A) Sympathetic reflex (ΔCSI) does not correlate with baseline blood pressure. (B) ΔCSI during the sit-to-stand test do not relate to concurrent changes in blood pressure during the protocol. (C) A significant negative correlation is found between resting CSI and baseline heart rate in controls, but not in stroke patients. (D) ΔCSI is significantly correlated with changes in heart rate during the protocol in both groups.

### CO_2_ regulation and sympathetic responsiveness

In participants with altered CO₂ regulation, a strong negative association was found between baseline EtCO₂ and the amplitude of the CSI response to standing, in both controls and stroke patients (Figure 4A; Spearman Correlation, EtCO_2_ constrained p=0.0081, r=-0.6299, EtCO_2_ unconstrained p=0.3467, r=-0.1523). Note that the definition of constrained ventilatory status was set at EtCO_2_ <35 mmHg, based on the literature [46–48], and confirmed in a data-driven approach as the threshold maximizing the relationship between the CSI response and EtCO_2_ in the constrained group (see Supplementary Figure 1). This relationship was not driven by a specific stroke hemisphere (Figures 4B and 4C), indicating that the link between CO_2_ regulation and sympathetic reflex is not necessarily driven by having had a stroke and its lateralization.

**Figure 4.**
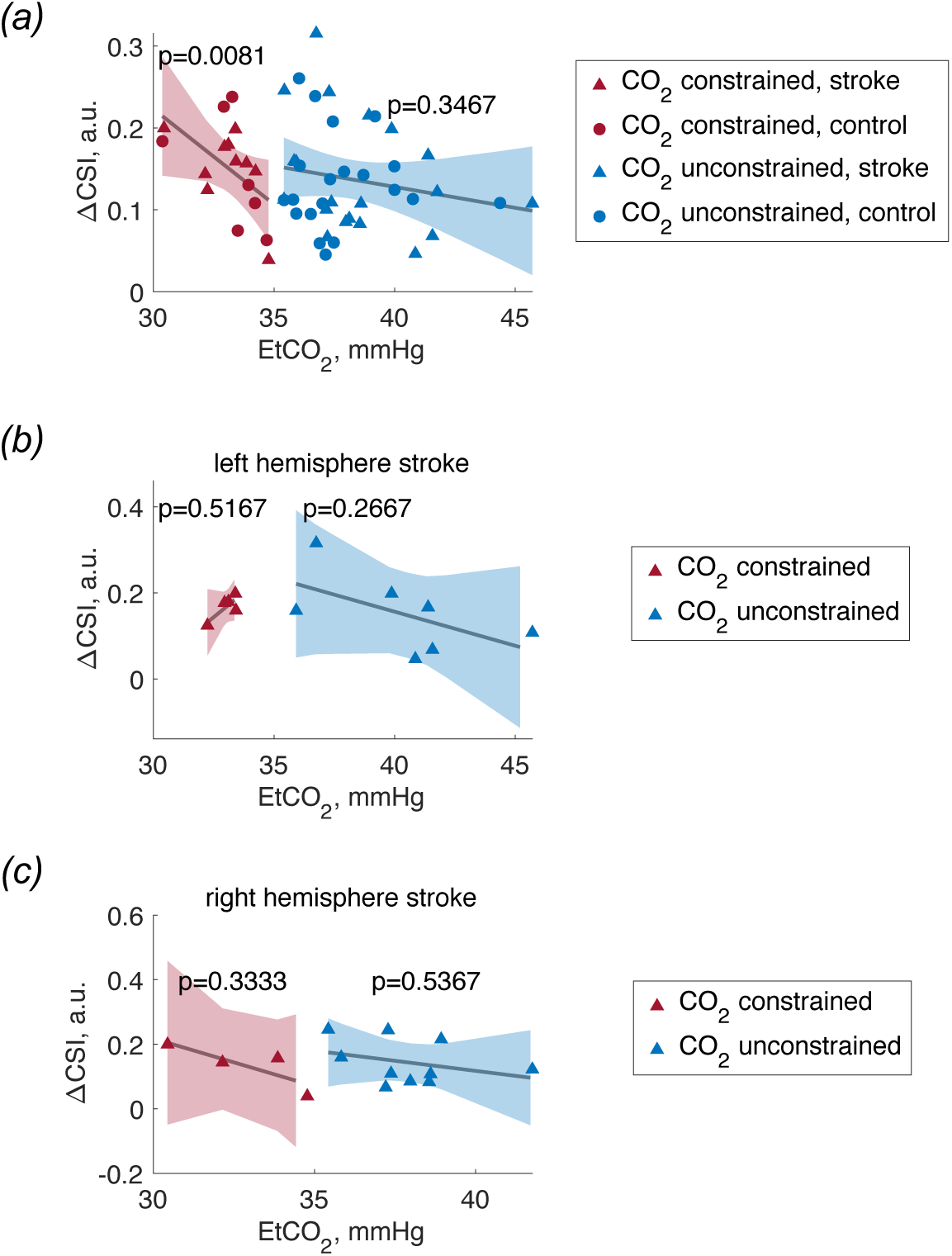
Influence of CO₂ regulation on the cardiac sympathetic reflex. (A) A negative correlation is observed between baseline end-tidal CO_2_ (EtCO_2_) levels and the magnitude of the CSI response to orthostatic stress in participants with constrained EtCO_2_, in both controls (n=7) and stroke patients (n=10). Relationship not found in unconstrained EtCO_2_ (controls, n=20; stroke patients, n=20). (B, C) This relationship is not driven by stroke hemisphere, indicating that CO_2_-related modulation of sympathetic activity is preserved across the whole subgroup (constrained EtCO_2_, left stroke, n=5, right stroke, n=4; unconstrained EtCO_2_, left stroke, n=7, right stroke, n=10).

Further analyses confirmed the specificity of this relationship. EtCO*_2_* values did not correlate with the changes in blood pressure (Figure 5A; Spearman Correlation, EtCO_2_ constrained p=0.7241, r=0.1, EtCO_2_ unconstrained p=0.2095, r=0.2139) or heart rate (Figure 5B; Spearman Correlation, EtCO_2_ constrained p=0.7144, r=-0.1036, EtCO_2_ unconstrained p=0.8007, r=-0.0435) during the orthostatic challenge, reinforcing the idea that the observed CO_2_–cardiac sympathetic association is not captured by conventional cardiovascular measures.

**Figure 5.**
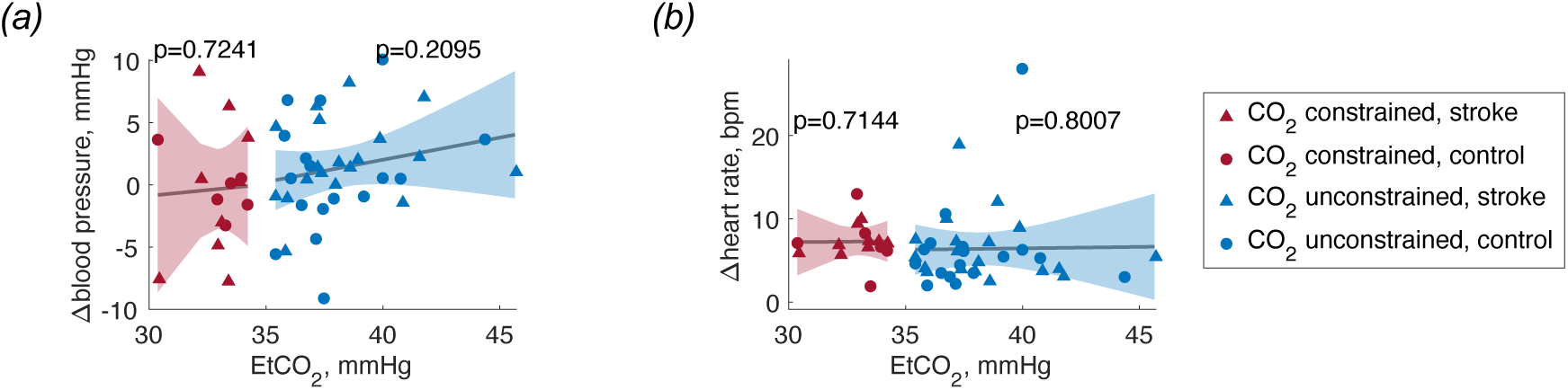
Specificity of the baseline end-tidal CO_2_ (EtCO_2_)–CSI relationship. No significant correlations are found between EtCO_2_ and changes in blood pressure (A) or heart rate (B) during the sit-to-stand test (constrained EtCO_2_, controls, n=7, stroke patients, n=10; unconstrained EtCO_2_, controls, n=20, stroke patients, n=20).

### Cerebral hemodynamics and sympathetic drive

Finally, we explored whether the CSI response to standing was associated with changes in cerebral blood flow. CSI correlated significantly with the percentage change in cerebral blood flow velocity in the right middle cerebral artery, from baseline to both sitting and standing phases (Figures 6A). A stronger correlation was found in the case of baseline vs sit (Spearman correlation, baseline vs sit p=0.0009, r=0.6239, baseline vs stand p=0.0365, r=0.4140). This effect was not observed in the case of the left middle cerebral artery. Notably, this coupling between sympathetic activity and the right middle cerebral artery hemodynamics, during the baseline to sit change, was preserved in stroke patients, without a tendency in one or another hemisphere of the stroke (Figure 6B; Spearman correlation p=0.0072, r=0.6559). An anticorrelated effect was found when performing the same analysis over the radial arteries (see Supplementary Figure 4).

**Figure 6.**
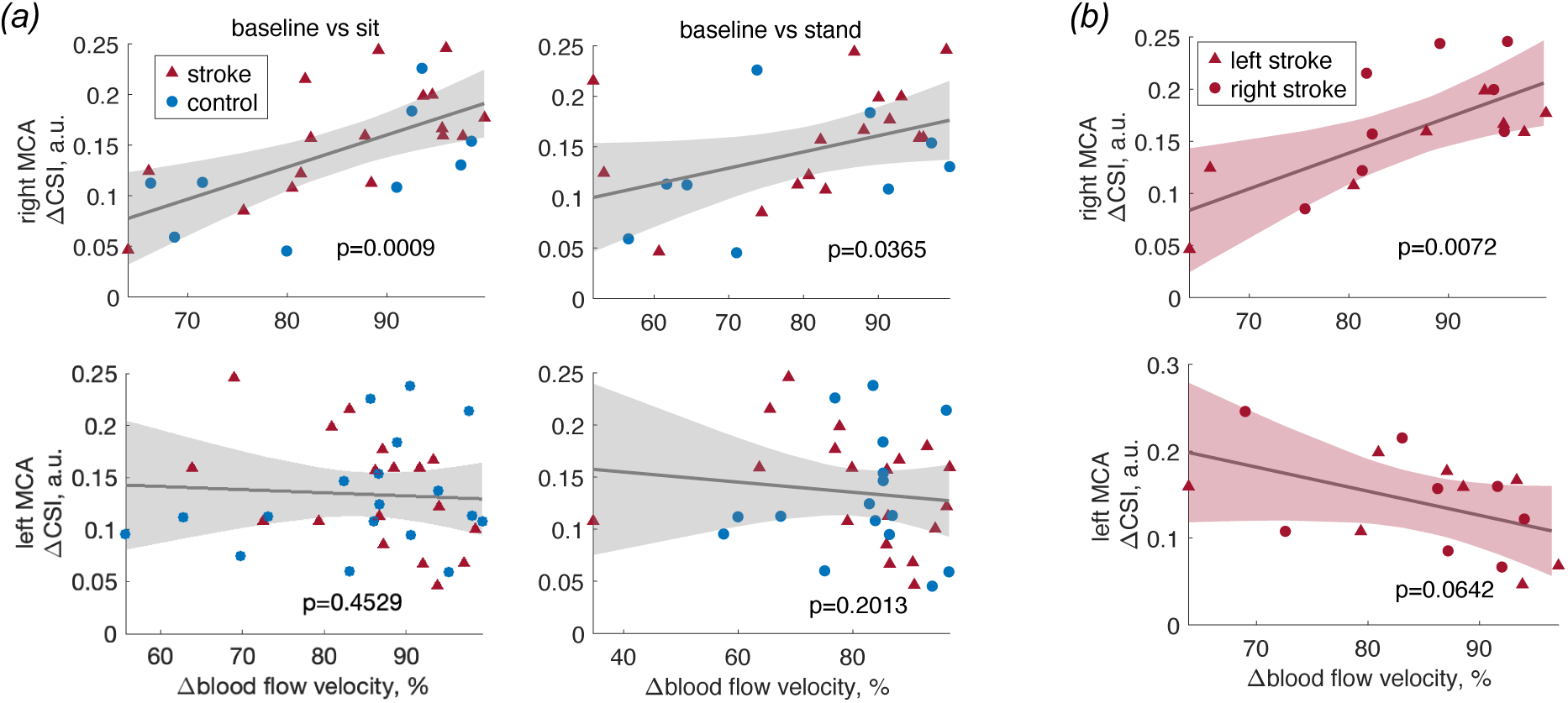
Association between sympathetic reflex and cerebral blood flow in the middle cerebral artery (MCA; right hemisphere, left hemisphere). (A) CSI changes during sit-to-stand correlate, only in the right MCA, with the percentage change in cerebral blood flow velocity, both from baseline to sitting and from baseline to standing. (B) This association is preserved in stroke patients. Right MCA, control, n=9, left stroke, n=8, right stroke, n=8, stroke no hemisphere information, n=1. Left MCA, control, n=18, left stroke, n=8, right stroke, n=8, stroke no hemisphere information, n=2.

In contrast, no such relationships were found when analysing changes in blood pressure (Figure 7A; Spearman correlation p=0.5123, r=-0.1369) or heart rate alone (Figure 7B; Spearman correlation p=0.0575, r=0.3862), underscoring the unique role of CSI in capturing relevant cardiac autonomic-cerebral interactions.

**Figure 7.**
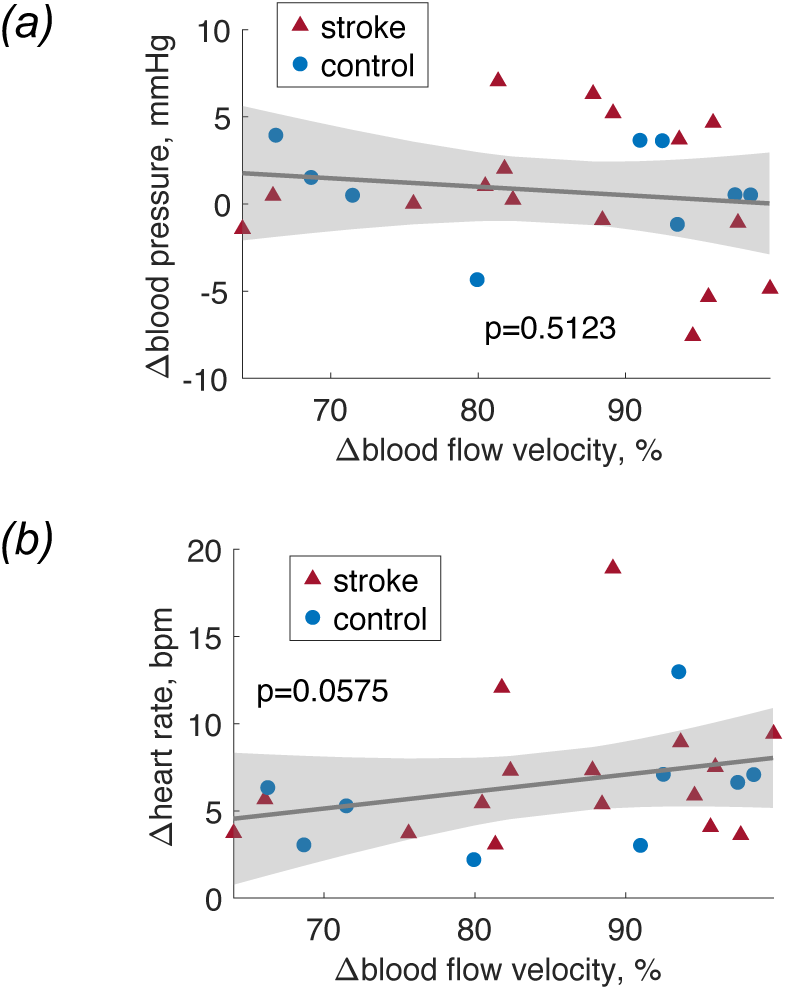
Lack of association between cerebral blood flow changes in the right middle cerebral artery and conventional cardiovascular markers. Changes in blood pressure (A) and heart rate (B) during the sit-to-stand test do not correlate with changes in cerebral blood flow velocity. Control, n=9, stroke, n=17.

## Discussion

This study provides compelling evidence that cardiac sympathetic reflexes under orthostatic stress provide meaningful information about potentially impaired cerebrovascular mechanisms. The relationships were related to cerebral autoregulation, which is affected not only by the presence of cerebrovascular or autonomic disease but also by the underlying ventilatory status, particularly CO_2_ regulation. In a cohort of older adults, including stroke survivors and matched controls, we found that while cardiac sympathetic reflexes in response to orthostatic stress were broadly preserved across groups, although marginally smaller in the stroke group, some minimal differences were indicative of maladaptive cerebral hemodynamics (blood flow velocity) and constrained ventilatory status (EtCO_2_).

Cerebral autoregulation emerges from the combined influence of partially overlapping control mechanisms. Small resistance vessels mediate myogenic adjustments to changes in arterial pressure; metabolic and chemical mechanisms can affect vascular tone; and mechanical factors such as intracranial pressure fluctuations further constrain flow [9,53,54]. Although the autonomic nervous system shapes the systemic cardiovascular inputs reaching the cerebral circulation, its direct role in autoregulatory control remains debated [4,5]. Rather than acting as a primary driver of autoregulation, autonomic activity likely interacts with these mechanisms indirectly by altering cardiac output and perfusion pressure [4], being these mechanisms actively involved under orthostatic stress and associated pathology [55]. In this context, our interpretation focuses on the interdependent regulatory loops linking autonomic responses, ventilatory CO_2_ control, and downstream microvascular adjustments, without assuming direct neural governance of autoregulatory behaviour.

The link between heightened sympathetic reactivity and exaggerated MCA velocity shifts may reflect impaired dynamic autoregulation, where arterioles fail to adequately counter sudden drops in perfusion pressure during orthostasis [9,54]. In such cases, systemic sympathetic responses may exceed, producing disproportionate pressure-flow transmission into the cerebral circulation. This maladaptive coupling aligns with models suggesting that autonomic overactivation emerges when cerebrovascular buffering is reduced, leading to a reliance on upstream cardiac responses to stabilize perfusion [56].

Our results show that elevated cardiac sympathetic responsiveness during a sit-to-stand test was specifically associated with lower resting EtCO_2_ levels, but only in patients with evidence of CO_2_ constraint. This relationship was not observed in participants with normal EtCO_2_, regardless of stroke status. We acknowledge that the literature on CO_2_ involvement in cerebrovascular coupling is mixed, particularly in humans compared to animal models, and that neural control pathways can modulate cerebral blood flow independently of CO_2_. Indeed, neurovascular coupling appears unconnected to CO_2_ as they relate to distinct regulations [57]. We found that all patients had a percentual change in blood flow velocity in the abnormal range (>10%), and that change scaled with cardiac sympathetic responses to orthostatic stress as well. These results suggest that increased sympathetic reactivity is associated with impaired cerebral hemodynamics and ventilatory inefficiency and may reflect patterns consistent with compensatory or maladaptive responses, although no causal relationship can be inferred from the present data.

Ventilatory status can exert an influence on cerebrovascular tone through pH-mediated changes in smooth muscle activity [58,59]. Even mild hypocapnia causes cerebral vasoconstriction and reduces the buffering capacity of the autoregulatory system. Low EtCO_2_ therefore imposes a state of reduced cerebrovascular reserve, making the circulation more dependent on autonomic compensation during postural stress [60]. In individuals with CO_2_ constraint, sympathetic activation during sit-to-stand may reflect an attempt to maintain perfusion in the setting of reduced cerebrovascular reserve, rather than a direct effect of CO_2_ on cerebrovascular regulation. This pattern is consistent with the exaggerated MCA velocity changes we observed.

While ventilatory inefficiency is traditionally attributed to impaired central respiratory drive or reduced pulmonary perfusion, low EtCO_2_ may also signify impaired cerebrovascular CO_2_ responsiveness, which we found to be present in participants with stroke as well. Importantly, this cardiac autonomic-ventilatory association was paralleled by exaggerated changes in cerebral blood flow velocity in the right middle cerebral artery, in response to postural shifts, which occurred in most participants. Heightened sympathetic activation exhibited larger increases in cerebral blood flow velocity, possibly indicating overcompensation due to impaired autoregulatory buffering. This finding aligns with the expected maladaptive neurovascular crosstalk, wherein autonomic and cerebrovascular systems meant to cooperate fail to maintain homeostasis when one component is compromised [6]. That the right middle central artery was predominantly affected may point to a lateralized role in integrating motor-postural and autonomic regulation, consistent with previous reports suggesting right-hemisphere dominance in autonomic control [42] and postural balance [43–45].

Ageing and stroke could further complicate these physiological interactions by altering arterial stiffness, baroreflex sensitivity, neurovascular coupling, and cerebrovascular CO_2_ responsiveness [61–63]. Structural vascular changes reduce compliance and impair the ability of microvessels to buffer pulsatile pressure, while stroke-related network disruption can degrade coordinated blood pressure and ventilatory responses. Thus, the maladaptive cardiac–cerebrovascular patterns observed may reflect the combined influence of physiological factors, including vascular stiffening, impaired CO₂ reactivity, and reduced baroreflex-mediated stability, even in individuals without overt stroke-related hemodynamic deficits.

While impaired cerebral autoregulation can be expected in chronic stroke patients [64], stroke history alone did not show distinct cardiac sympathetic reactivity, blood flow velocity or EtCO_2_ profiles in this cohort, suggesting that maladaptive patterns are not exclusively a result of focal lesions but may emerge from broader network-level disintegration and systemic vulnerabilities that occur with ageing [65,66]. Still, stroke may act as a sensitizing event that unmasks or exacerbates latent cardiac, vascular and respiratory inefficiencies [67], particularly when autoregulatory control is weakened. It is also possible that the heterogeneity of lesion locations and small sample size diluted group-level effects, further highlighting the need for stratified approaches based on functional phenotypes (e.g., CO_2_-constrained vs. non-constrained [68]) rather than clinical diagnosis alone.

Although CSI is derived from the Poincaré representation of RR-interval dynamics, in this study it was computed using a time-resolved framework and then condition-averaged [26], rather than obtained from a static Poincaré plot over the entire recording. Our focus on CSI reflects its role as a recently validated [26,41,69], sympathetic-specific index with demonstrated sensitivity to autonomic elicitation. However, we acknowledge that additional Poincaré-based descriptors [70,71] may provide complementary insight into autonomic modulation. A systematic comparison of these measures represents an important direction for future work but lies outside the scope of the present study, which was designed to evaluate CSI as a targeted marker of sympathetic reactivity in relation to cerebral autoregulation and ventilatory status.

Taken together, these mechanisms suggest that cardiac sympathetic responses during orthostatic stress do not act in isolation but emerge from the integrated behaviour of many physiological systems. Individuals with constrained CO_2_ regulation, and thus reduced cerebrovascular reserve, may rely more on sympathetic compensation to maintain perfusion, leading to exaggerated MCA velocity changes. Our findings therefore support a model in which ventilatory inefficiency diminishes cerebrovascular buffering capacity, and autonomic activation may fill the resulting gap [10,60]. This interaction provides a potential physiological substrate for the broader cardiac–cerebrovascular crosstalk observed in ageing and stroke.

We acknowledge limitations of our study. First, although EtCO_2_ is commonly used as an index of ventilatory status, it is not a direct measure of arterial CO_2_ (PaCO_2_). The EtCO_2_– PaCO_2_ relationship can be influenced by many factors including tidal volume, respiratory rate, ventilation-perfusion mismatch, and alterations in dead space [72]. Therefore, while EtCO_2_ is an accessible, non-invasive surrogate for relative changes in ventilatory regulation, it could be a biased estimator of arterial CO_2_. In addition, our EtCO_2_ measure was based on a single value rather than continuous reactivity curves, which may limit sensitivity. Second, although the sit-to-stand paradigm is ecologically valid, it lacks the precision of controlled tilt-table testing. Third, we did not assess dynamic baroreflex sensitivity or direct cerebral oxygenation, which could provide further granularity to the interplay between cardiovascular and cerebrovascular responses. Fourth, this study did not include measurements in the posterior cerebral arteries, which have shown certain specificity in capturing reduced perfusion in elderly under orthostatic stress [73]. However, it is important to note that the posterior cerebral artery may receive collaterals from the anterior circulation via the posterior communicating arteries and may therefore be a poor surrogate for the basilar artery, which is the main supply to the brain stem [74,75]. Unfortunately, simultaneous continuous monitoring of the basilar, posterior and middle cerebral arteries during the sit-to-stand protocol was not available. Finally, the present analytical approach is primarily based on correlation analyses. While this strategy is appropriate given the sample size, heterogeneity, and incomplete availability of physiological measures across participants, it inherently limits the scope of statistical inference. In particular, correlation-based analyses do not permit conclusions regarding causality and do not account for the simultaneous interactions among multiple physiological variables. A more comprehensive multivariate modelling framework would be desirable to address these complex interrelationships; however, the limited sample size and missing data across variables precluded the reliable application of such approaches in the current study. Accordingly, the findings should be interpreted as descriptive associations rather than evidence of causal mechanisms.

From a translational perspective, our findings emphasize the need to consider ventilatory efficiency and autonomic capacity jointly when assessing cerebrovascular risk in the elderly [68]. While autonomic testing is routine in some clinical settings, the incorporation of novel cardiac-based measures [26] under simple postural challenges could enhance our ability to detect subclinical dysregulations. Importantly, we demonstrated that cardiac sympathetic responses are better reflected in the CSI, as compared to standard measures [34], such as heart rate and blood pressure. The relevance of these findings highlight that cardiac sympathetic activity and cardiovascular coupling do not necessarily correlate [76], and may convey information about distinct mechanisms. Future research can further enlighten how brain-heart interactions are disrupted in the elderly and in stroke patients, helping to clarify the role of these dynamics in the regulatory mechanisms involved in postural control [77].

In conclusion, our findings support a model in which impaired cerebrovascular dynamics and ventilatory status modifies the expression of cardiac autonomic responses to postural stress in aged individuals. This cardiac-cerebrovascular crosstalk appears most pronounced in CO_2_-constrained individuals, regardless of stroke history, and may underlie a broader phenotype of age-related physiological decline. These insights contribute to the emerging framework of brain-heart interactions and underscore the importance of multidomain physiological profiling in ageing and neurovascular disease.

## Acknowledgements

Diego Candia-Rivera is supported by the European Commission, Horizon MSCA Postdoctoral Fellowship Program (grant n° 101151118).

## References

1. Candia-Rivera D, Faes L, Fallani FDV, Chavez M. 2025 Measures and Models of Brain-Heart Interactions. IEEE Reviews in Biomedical Engineering **in press**. (doi:10.1109/RBME.2025.3529363)

2. Phillips AA, Chan FH, Zheng MMZ, Krassioukov AV, Ainslie PN. 2016 Neurovascular coupling in humans: Physiology, methodological advances and clinical implications. J Cereb Blood Flow Metab 36, 647–664. (doi:10.1177/0271678X15617954)

3. Blinder P, Tsai PS, Kaufhold JP, Knutsen PM, Suhl H, Kleinfeld D. 2013 The cortical angiome: an interconnected vascular network with noncolumnar patterns of blood flow. Nat Neurosci 16, 889–897. (doi:10.1038/nn.3426)

4. Ainslie PN, Brassard P. 2014 Why is the neural control of cerebral autoregulation so controversial? F1000Prime Rep 6, 14. (doi:10.12703/P6-14)

5. Brassard P, Tymko MM, Ainslie PN. 2017 Sympathetic control of the brain circulation: Appreciating the complexities to better understand the controversy. Auton Neurosci 207, 37–47. (doi:10.1016/j.autneu.2017.05.003)

6. Aaslid R, Lindegaard KF, Sorteberg W, Nornes H. 1989 Cerebral autoregulation dynamics in humans. Stroke 20, 45–52. (doi:10.1161/01.STR.20.1.45)

7. Engelen T, Solcà M, Tallon-Baudry C. 2023 Interoceptive rhythms in the brain. Nat Neurosci 26, 1670–1684. (doi:10.1038/s41593-023-01425-1)

8. Tan CO, Taylor JA. 2014 Integrative physiological and computational approaches to understand autonomic control of cerebral autoregulation. Experimental Physiology 99, 3– 15. (doi:10.1113/expphysiol.2013.072355)

9. Claassen JA, Meel-van den Abeelen AS, Simpson DM, Panerai RB. 2016 Transfer function analysis of dynamic cerebral autoregulation: A white paper from the International Cerebral Autoregulation Research Network. J Cereb Blood Flow Metab 36, 665–680. (doi:10.1177/0271678X15626425)

10. Willie CK, Tzeng Y-C, Fisher JA, Ainslie PN. 2014 Integrative regulation of human brain blood flow. J Physiol 592, 841–859. (doi:10.1113/jphysiol.2013.268953)

11. Zhang R, Zuckerman JH, Iwasaki K, Wilson TE, Crandall CG, Levine BD. 2002 Autonomic neural control of dynamic cerebral autoregulation in humans. Circulation 106, 1814–1820. (doi:10.1161/01.cir.0000031798.07790.fe)

12. Ogoh S, Brothers RM, Eubank WL, Raven PB. 2008 Autonomic Neural Control of the Cerebral Vasculature. Stroke 39, 1979–1987. (doi:10.1161/STROKEAHA.107.510008)

13. Hamner JW, Tan CO, Lee K, Cohen MA, Taylor JA. 2010 Sympathetic Control of the Cerebral Vasculature in Humans. Stroke 41, 102–109. (doi:10.1161/STROKEAHA.109.557132)

14. Chen HS, van Roon L, Ge Y, van Gils JM, Schoones JW, DeRuiter MC, Zeppenfeld K, Jongbloed MRM. 2024 The relevance of the superior cervical ganglion for cardiac autonomic innervation in health and disease: a systematic review. Clin Auton Res 34, 45– 77. (doi:10.1007/s10286-024-01019-2)

15. Attwell D, Buchan AM, Charpak S, Lauritzen M, MacVicar BA, Newman EA. 2010 Glial and neuronal control of brain blood flow. Nature 468, 232–243. (doi:10.1038/nature09613)

16. Iadecola C. 2017 The Neurovascular Unit Coming of Age: A Journey through Neurovascular Coupling in Health and Disease. Neuron 96, 17–42. (doi:10.1016/j.neuron.2017.07.030)

17. DiNuzzo M et al. 2024 Neurovascular coupling is optimized to compensate for the increase in proton production from nonoxidative glycolysis and glycogenolysis during brain activation and maintain homeostasis of pH, pCO2, and pO2. J Neurochem 168, 632–662. (doi:10.1111/jnc.15839)

18. Panerai RB, Deverson ST, Mahony P, Hayes P, Evans DH. 1999 Effect of CO2 on dynamic cerebral autoregulation measurement. Physiol. Meas. 20, 265. (doi:10.1088/0967-3334/20/3/304)

19. Wheeler C, Pacheco JM, Kim AC, Camacho-Santiago M, Kalafut MA, Ahern T, White AA, Patay B, Criado JR. 2022 Cardiovascular Autonomic Regulation, ETCO2 and the Heart Rate Response to the Tilt Table Test in Patients with Orthostatic Intolerance. Appl Psychophysiol Biofeedback 47, 107–119. (doi:10.1007/s10484-022-09536-4)

20. Burzyńska M, Woźniak J, Urbański P, Kędziora J, Załuski R, Goździk W, Uryga A. 2025 Heart Rate Variability and Cerebral Autoregulation in Patients with Traumatic Brain Injury with Paroxysmal Sympathetic Hyperactivity Syndrome. Neurocrit Care 42, 864–877. (doi:10.1007/s12028-024-02149-1)

21. Head GA. 1995 Baroreflexes and Cardiovascular Regulation in Hypertension. Journal of Cardiovascular Pharmacology 26, S7.

22. Xing C-Y et al. 2017 Arterial Pressure, Heart Rate, and Cerebral Hemodynamics Across the Adult Life Span. Hypertension 69, 712–720. (doi:10.1161/HYPERTENSIONAHA.116.08986)

23. Wang S, Wang X, Zhao Y, Xie L, Zhang J. 2025 A feedback loop study of brain-heart interaction based on HEP and HRV. Biocybernetics and Biomedical Engineering 45, 181–188. (doi:10.1016/j.bbe.2025.02.005)

24. Doehner W et al. 2018 Heart and brain interaction in patients with heart failure: overview and proposal for a taxonomy. A position paper from the Study Group on Heart and Brain Interaction of the Heart Failure Association. European Journal of Heart Failure 20, 199– 215. (doi:10.1002/ejhf.1100)

25. Olufsen MS, Tran HT, Ottesen JT, Research Experiences for Undergraduates Program, Lipsitz LA, Novak V. 2006 Modeling baroreflex regulation of heart rate during orthostatic stress. Am J Physiol Regul Integr Comp Physiol 291, R1355–1368. (doi:10.1152/ajpregu.00205.2006)

26. Candia-Rivera D, de Vico Fallani F, Chavez M. 2025 Robust and time-resolved estimation of cardiac sympathetic and parasympathetic indices. Royal Society Open Science 12, 240750. (doi:10.1098/rsos.240750)

27. Cheshire WP, Goldstein DS. 2019 Autonomic uprising: the tilt table test in autonomic medicine. Clin Auton Res 29, 215–230. (doi:10.1007/s10286-019-00598-9)

28. Reyes del Paso GA, Langewitz W, Mulder LJM, van Roon A, Duschek S. 2013 The utility of low frequency heart rate variability as an index of sympathetic cardiac tone: a review with emphasis on a reanalysis of previous studies. Psychophysiology 50, 477–487. (doi:10.1111/psyp.12027)

29. Castro P, Serrador J, Sorond F, Azevedo E, Rocha I. 2022 Sympathovagal imbalance in early ischemic stroke is linked to impaired cerebral autoregulation and increased infarct volumes. Autonomic Neuroscience 241, 102986. (doi:10.1016/j.autneu.2022.102986)

30. Lakatos LB, Shin DC, Müller M, Österreich M, Marmarelis V, Bolognese M. 2024 Impaired dynamic cerebral autoregulation measured in the middle cerebral artery in patients with vertebrobasilar ischemia is associated with autonomic failure. Journal of Stroke and Cerebrovascular Diseases 33, 107454. (doi:10.1016/j.jstrokecerebrovasdis.2023.107454)

31. Nasr N, Czosnyka M, Pavy-Le Traon A, Custaud M-A, Liu X, Varsos GV, Larrue V. 2014 Baroreflex and Cerebral Autoregulation Are Inversely Correlated. Circulation Journal 78, 2460–2467. (doi:10.1253/circj.CJ-14-0445)

32. Korpelainen JT, Sotaniemi KA, Myllylä VV. 1999 Autonomic nervous system disorders in stroke. Clinical Autonomic Research 9, 325–333. (doi:10.1007/BF02318379)

33. Goldberger AL et al. 2000 PhysioBank, PhysioToolkit, and PhysioNet: components of a new research resource for complex physiologic signals. Circulation 101, E215–220. (doi:10.1161/01.cir.101.23.e215)

34. Novak V, Hu K, Desrochers L, Novak P, Caplan L, Lipsitz L, Selim M. 2010 Cerebral flow velocities during daily activities depend on blood pressure in patients with chronic ischemic infarctions. Stroke 41, 61–66. (doi:10.1161/STROKEAHA.109.565556)

35. Low PA, Opfer-Gehrking TL, Proper CJ, Zimmerman I. 1990 The effect of aging on cardiac autonomic and postganglionic sudomotor function. Muscle Nerve 13, 152–157. (doi:10.1002/mus.880130212)

36. 1996 Consensus statement on the definition of orthostatic hypotension, pure autonomic failure, and multiple system atrophy. The Consensus Committee of the American Autonomic Society and the American Academy of Neurology. Neurology 46, 1470. (doi:10.1212/wnl.46.5.1470)

37. Pan J, Tompkins WJ. 1985 A Real-Time QRS Detection Algorithm. IEEE Transactions on Biomedical Engineering 32, 230–236. (doi:10.1109/TBME.1985.325532)

38. Candia-Rivera D, Catrambone V, Barbieri R, Valenza G. 2021 Integral pulse frequency modulation model driven by sympathovagal dynamics: Synthetic vs. real heart rate variability. Biomedical Signal Processing and Control 68, 102736. (doi:10.1016/j.bspc.2021.102736)

39. Candia-Rivera D, Chavez M. 2025 Cardiac-vagal rhythm echoes on the heartbeat’s mechanosensory imprint in the brain. Commun Biol 8, 1578. (doi:10.1038/s42003-025-08969-x)

40. Candia-Rivera D, Chavez M. 2024 Tracking the physiological responses in sleep apnea using robust cardiac sympathetic activity estimation. In 2024 13th Conference of the European Study Group on Cardiovascular Oscillations (ESGCO), pp. 1–2. (doi:10.1109/ESGCO63003.2024.10766963)

41 Candia-Rivera D, Carrion-Falgarona S, Fallani F de V, Chavez M. 2025 Modeling the time-resolved modulations of cardiac activity in rats: A study on pharmacological autonomic stimulation. Journal of Physiology in press. (doi:10.1113/JP288400)

42. Hilz MJ, Dütsch M, Perrine K, Nelson PK, Rauhut U, Devinsky O. 2001 Hemispheric influence on autonomic modulation and baroreflex sensitivity. Annals of Neurology 49, 575–584. (doi:10.1002/ana.1006)

43. Manor B, Hu K, Zhao P, Selim M, Alsop D, Novak P, Lipsitz L, Novak V. 2010 Altered control of postural sway following cerebral infarction. Neurology 74, 458–464. (doi:10.1212/WNL.0b013e3181cef647)

44. Pérennou D, Pélissier J, Amblard B. 1996 La posture et le contrôle postural du patient cérébrolésé vasculaire: une revue de la littérature. Annales de Réadaptation et de Médecine Physique 39, 497–513. (doi:10.1016/S0168-6054(97)84233-X)

45. Spinazzola L, Cubelli R, Della Sala S. 2003 Impairments of trunk movements following left or right hemisphere lesions: dissociation between apraxic errors and postural instability. Brain 126, 2656–2666. (doi:10.1093/brain/awg266)

46. Dong L, Takeda C, Kamitani T, Hamada M, Hirotsu A, Yamamoto Y, Mizota T. 2023 Association between intraoperative end-tidal carbon dioxide and postoperative organ dysfunction in major abdominal surgery: A cohort study. PLoS One 18, e0268362. (doi:10.1371/journal.pone.0268362)

47. Dong L, Takeda C, Yamazaki H, Kamitani T, Kimachi M, Hamada M, Fukuhara S, Mizota T, Yamamoto Y. 2021 Intraoperative end-tidal carbon dioxide and postoperative mortality in major abdominal surgery: a historical cohort study. Can J Anesth/J Can Anesth 68, 1601–1610. (doi:10.1007/s12630-021-02086-z)

48. Le Gall A, Eustache G, Coquet A, Seguin P, Launey Y. 2024 End-tidal carbon dioxide and arterial to end-tidal carbon dioxide gradient are associated with mortality in patients with neurological injuries. Sci Rep 14, 19172. (doi:10.1038/s41598-024-69143-7)

49. Garrett ZK, Pearson J, Subudhi AW. 2017 Postural effects on cerebral blood flow and autoregulation. Physiol Rep 5, e13150. (doi:10.14814/phy2.13150)

50. Aries MJ, Elting JW, Stewart R, De Keyser J, Kremer B, Vroomen P. 2013 Cerebral blood flow velocity changes during upright positioning in bed after acute stroke: an observational study. BMJ Open 3, e002960. (doi:10.1136/bmjopen-2013-002960)

51. Ho J, Tumkaya T, Aryal S, Choi H, Claridge-Chang A. 2019 Moving beyond P values: data analysis with estimation graphics. Nat Methods 16, 565–566. (doi:10.1038/s41592-019-0470-3)

52. Vargha A, Delaney HD. 2000 A critique and improvement of the CL common language effect size statistics of McGraw and Wong. Journal of Educational and Behavioral Statistics 25, 101–132. (doi:10.2307/1165329)

53. Meng L, Gelb AW. 2015 Regulation of cerebral autoregulation by carbon dioxide. Anesthesiology 122, 196–205. (doi:10.1097/ALN.0000000000000506)

54. Tzeng Y-C, Ainslie PN. 2014 Blood pressure regulation IX: cerebral autoregulation under blood pressure challenges. Eur J Appl Physiol 114, 545–559. (doi:10.1007/s00421-013-2667-y)

55. Furlan R, Barbic F, Casella F, Severgnini G, Zenoni L, Mercieri A, Mangili R, Costantino G, Porta A. 2009 Neural autonomic control in orthostatic intolerance. Respiratory Physiology & Neurobiology 169, S17–S20. (doi:10.1016/j.resp.2009.04.008)

56. Olufsen MS, Ottesen JT, Tran HT, Ellwein LM, Lipsitz LA, Novak V. 2005 Blood pressure and blood flow variation during postural change from sitting to standing: model development and validation. J Appl Physiol (1985) 99, 1523–1537. (doi:10.1152/japplphysiol.00177.2005)

57. Tournissac M et al. 2024 Neurovascular coupling and CO2 interrogate distinct vascular regulations. Nat Commun 15, 7635. (doi:10.1038/s41467-024-49698-9)

58. Lucas SJE et al. 2011 Alterations in cerebral blood flow and cerebrovascular reactivity during 14 days at 5050 m. The Journal of Physiology 589, 741–753. (doi:10.1113/jphysiol.2010.192534)

59. Battisti-Charbonney A, Fisher J, Duffin J. 2011 The cerebrovascular response to carbon dioxide in humans. The Journal of Physiology 589, 3039–3048. (doi:10.1113/jphysiol.2011.206052)

60. Ainslie PN, Duffin J. 2009 Integration of cerebrovascular CO2 reactivity and chemoreflex control of breathing: mechanisms of regulation, measurement, and interpretation. *American Journal of Physiology-Regulatory*, Integrative and Comparative Physiology 296, R1473–R1495. (doi:10.1152/ajpregu.91008.2008)

61. Aries MJH, Elting JW, De Keyser J, Kremer BPH, Vroomen PCAJ. 2010 Cerebral Autoregulation in Stroke. Stroke 41, 2697–2704. (doi:10.1161/STROKEAHA.110.594168)

62. Marín J, Rodríguez-Martínez MA. 1999 Age-related changes in vascular responses. Experimental Gerontology 34, 503–512. (doi:10.1016/S0531-5565(99)00029-7)

63. Tarumi T, Ayaz Khan M, Liu J, Tseng BY, Parker R, Riley J, Tinajero C, Zhang R. 2014 Cerebral hemodynamics in normal aging: central artery stiffness, wave reflection, and pressure pulsatility. J Cereb Blood Flow Metab 34, 971–978. (doi:10.1038/jcbfm.2014.44)

64. Aoi MC, Hu K, Lo M-T, Selim M, Olufsen MS, Novak V. 2012 Impaired cerebral autoregulation is associated with brain atrophy and worse functional status in chronic ischemic stroke. PLoS One 7, e46794. (doi:10.1371/journal.pone.0046794)

65. Kumral D et al. 2019 The age-dependent relationship between resting heart rate variability and functional brain connectivity. NeuroImage 185, 521–533. (doi:10.1016/j.neuroimage.2018.10.027)

66. Satoh K, Ohashi A, Kumagai M, Sato M, Kuji A, Joh S. 2015 Evaluation of Differences between PaCO2 and ETCO2 by Age as Measured during General Anesthesia with Patients in a Supine Position. Journal of Anesthesiology 2015, 710537. (doi:10.1155/2015/710537)

67. Smith AC, Saunders DH, Mead G. 2012 Cardiorespiratory Fitness after Stroke: A Systematic Review. International Journal of Stroke 7, 499–510. (doi:10.1111/j.1747-4949.2012.00791.x)

68. Dempsey JA, Smith CA. 2014 Pathophysiology of human ventilatory control. European Respiratory Journal 44, 495–512. (doi:10.1183/09031936.00048514)

69. Candia-Rivera D, Boehme R, Salamone PC. 2025 Autonomic modulations to cardiac dynamics in response to affective touch: Differences between social touch and self-touch. IEEE Transactions on Affective Computing, 1–11. (doi:10.1109/TAFFC.2025.3548778)

70. Porta A et al. 2008 Temporal asymmetries of short-term heart period variability are linked to autonomic regulation. *American Journal of Physiology-Regulatory*, Integrative and Comparative Physiology 295, R550–R557. (doi:10.1152/ajpregu.00129.2008)

71. Guzik P, Piskorski J, Krauze T, Wykretowicz A, Wysocki H. 2006 Heart rate asymmetry by Poincaré plots of RR intervals. Biomed Tech (Berl) 51, 272–275. (doi:10.1515/BMT.2006.054)

72. Jones NL, Robertson DG, Kane JW. 1979 Difference between end-tidal and arterial PCO2 in exercise. J Appl Physiol Respir Environ Exerc Physiol 47, 954–960. (doi:10.1152/jappl.1979.47.5.954)

73. Sorond FA, Khavari R, Serrador JM, Lipsitz LA. 2005 Regional Cerebral Autoregulation During Orthostatic Stress: Age-Related Differences. The Journals of Gerontology: Series A 60, 1484–1487. (doi:10.1093/gerona/60.11.1484)

74. Tatu L, Moulin T, Bogousslavsky J, Duvernoy H. 1996 Arterial territories of human brain. Neurology 47, 1125–1135. (doi:10.1212/WNL.47.5.1125)

75. Fujii K, Heistad DD, Faraci FM. 1991 Role of the basilar artery in regulation of blood flow to the brain stem in rats. Stroke 22, 763–767. (doi:10.1161/01.STR.22.6.763)

76. Xie L, Liu B, Wang X, Mei M, Li M, Yu X, Zhang J. 2017 Effects of different stresses on cardiac autonomic control and cardiovascular coupling. Journal of Applied Physiology 122, 435–445. (doi:10.1152/japplphysiol.00245.2016)

77. Dohata M, Kaneko N, Takahashi R, Suzuki Y, Nakazawa K. 2025 Posture-Dependent Modulation of Interoceptive Processing in Young Male Participants: A Heartbeat-Evoked Potential Study. European Journal of Neuroscience 61, e70021. (doi:10.1111/ejn.70021)

