## Supplementary material for "Cardiac-cerebrovascular crosstalk: Cardiac rhythms reveal maladaptive cerebral blood flow velocity and constrained ventilatory status"

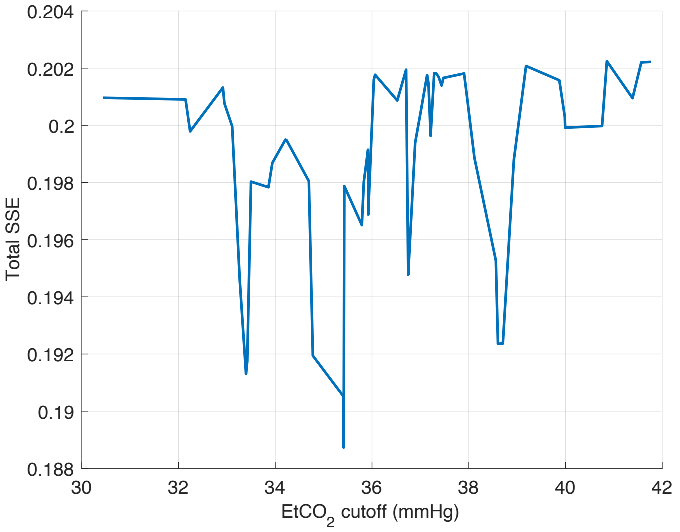


*Supplementary Figure 1. Influence of $EtCO_2$ threshold defining constrained ventilatory status in finding relationships between the cardiac sympathetic response to postural changes and the baseline $EtCO_2$ The minimum sum of squared errors (SSE) of the two correlations performed in the low and high EtCO2 groups indicates the optimal, data-driven threshold maximising the relationship.*

*Supplementary Table 1. Stroke aetiologies, hemispheres and regions affected. "-" indicates information unavailable.*

| id | Aetiology | Region | |
|---|---|---|---|
| 185 | Atherosclerosis (large vessel, intracranial) | L | Basal ganglia |
| 205 | Atherosclerosis (large vessel, intracranial) | R | Posterior MCA |
| 231 | - | L | Temporal |
| 232 | Atherosclerosis (large vessel, intracranial) | L | Posterior MCA |
| 239 | Atherosclerosis (large vessel, intracranial) | - | - |
| 240 | Atherosclerosis (large vessel, intracranial) | L | Frontoparietal |
| 244 | Atherosclerosis (large vessel, intracranial) | R | Temporal |
| 247 | Atherosclerosis (large vessel, intracranial) | R | Temporoparietal |
| 248 | Atherosclerosis (large vessel, intracranial) | L | Temporal |
| 261 | Atherosclerosis (large vessel, intracranial) | R | Internal capsule, basal ganglia |
| 277 | Atherosclerosis (large vessel, intracranial) | R | Temporal |
| 295 | Atherosclerosis (large vessel, intracranial) | R | Temporal, insular |
| 321 | Cardioembolism (hypercoagulable state) | L | Frontal pre-and postcentral gyrus |
| 322 | - | L , R | L: medial temporal, external capsule<br>R: cerebellum, thalamus lacunae |
| 332 | - | L | Temporal-parietal |
| 334 | - | - | - |
| 336 | Cardioembolism (Deep Vein Thrombosis - Patent Foramen Ovale) | R | - |
| 337 | Cardioembolism | R | Temporo-parietal, occipital |
| 351 | Atherothrombosis (large vessel) | L | Temporal |
| 353 | - | R | multiple site MCA |
| 354 | - | - | - |
| 358 | - | L | Temporal |
| 361 | Arterioembolic | - | - |
| 363 | Atherosclerosis (large vessel, intracranial) | R | Frontal, temporal |
| 371 | Atherothrombosis | L | Frontal, occipital |
| 374 | Atherosclerosis | L | Frontal |
| 379 | Atherothrombosis (aortic aneurysm dissection) | L, R | Frontal, basal ganglia |
| 388 | Atherosclerosis | L | L parietal, L temporal, L frontal infarcts |
| 389 | - | - | |
| 402 | Embolism (Patent Foramen Ovale) | L, R | Frontal |

*Supplementary Table 2. Hypertensive patient sub-cohort and medication use. Patients do not displaying any medication use have history of hypertension but were not treated against hypertension during the study. "-" indicates information unavailable.*

| id | Patient group | Years with hypertension | Hypertensive medication use | | | | |
|---|---|---|---|---|---|---|---|
| | | | Angiotensin-converting-enzyme inhibitors | Angiotensin II Receptor Blockers | Beta blockers | Diuretics | Calcium channel blockers |
| 64 | Control | - | | | | | |
| 164 | Control | - | | | | | |
| 165 | Control | 4 | | | | X | |
| 172 | Control | - | | | | | |
| 184 | Control | - | | | | | |
| 185 | Stroke | 3 | | X | | | |
| 194 | Control | - | | | | | |
| 200 | Control | - | | | | | |
| 203 | Control | 8 | X | | | | |
| 204 | Control | 6 | | | | X | X |
| 205 | Stroke | - | X | | | | |
| 207 | Control | 10 | | | | X | X |
| 208 | Control | - | | | | | |
| 212 | Control | 3 | X | | | X | |
| 213 | Control | - | | | | | |
| 215 | Control | 3 | X | | | | |
| 218 | Control | - | | | | | |
| 221 | Control | - | | | | | |
| 225 | Control | - | | | | | |
| 227 | Control | 16 | X | | | | |
| 228 | Control | - | | | | | |
| 231 | Stroke | 6 | | X | | | |
| 232 | Stroke | 6 | | | X | | |
| 239 | Stroke | 21 | X | | | | |
| 240 | Stroke | - | X | | | | |
| 242 | Control | 25 | | | | X | X |
| 243 | Control | 26 | X | | X | X | |
| 244 | Stroke | - | | | | | |
| 246 | Control | 31 | | | X | | X |
| 247 | Stroke | 1 | | | | X | |
| 248 | Stroke | 2 | X | | | X | |
| 261 | Stroke | - | | | | | |
| 277 | Stroke | 24 | X | | X | X | |
| 295 | Stroke | 4 | | X | | X | X |
| 305 | Control | - | | | | | |

| | | | | | | | |
|---|---|---|---|---|---|---|---|
| 321 | Stroke | - | | | | X | X |
| 322 | Stroke | 20 | | X | | | |
| 332 | Stroke | 1 | X | | | | |
| 334 | Stroke | - | | | | | |
| 336 | Stroke | - | | | | | |
| 337 | Stroke | - | | | | | |
| 343 | Control | 1 | X | | | | |
| 351 | Stroke | 12 | X | | | | X |
| 353 | Stroke | 4 | | | X | | |
| 354 | Stroke | - | | | | | |
| 358 | Stroke | 7 | | | | | |
| 361 | Stroke | 36 | X | | X | X | |
| 363 | Stroke | - | | | X | | X |
| 364 | Control | - | | | | | |
| 371 | Stroke | 4 | X | | | | |
| 374 | Stroke | 3 | X | | | | |
| 376 | Control | 4 | | | | X | |
| 379 | Stroke | 1 | X | | X | X | |
| 388 | Stroke | 1 | X | | | | |
| 389 | Stroke | - | | X | X | X | |
| 399 | Control | - | | | | | |
| 402 | Stroke | - | | | | | |

*Supplementary Table 3. Clinical scores of stroke patients, based on the National Institutes of Health Stroke Scale (NIHSS) and the modified Rankin scale (MRS). “-“ indicates information unavailable.*

| id | NIHSS | MRS |
|---|---|---|
| 185 | 0 | 0 |
| 205 | 10 | 3 |
| 231 | 1 | 0 |
| 232 | 1 | 1 |
| 239 | 8 | 2 |
| 240 | 3 | - |
| 244 | 1 | 1 |
| 247 | 0 | 0 |
| 248 | 1 | 0 |
| 261 | 7 | 2 |
| 277 | 0 | 0 |
| 295 | 4 | 3 |
| 321 | 1 | 1 |
| 322 | 2 | 3 |
| 332 | 1 | 0 |
| 334 | 2 | 2 |
| 336 | 0 | 0 |
| 337 | 3 | 1 |
| 351 | 1 | 1 |
| 353 | 2 | 1 |
| 354 | 2 | 1 |
| 358 | 1 | 0 |
| 361 | 8 | 3 |
| 363 | 6 | 3 |
| 371 | 3 | 1 |
| 374 | 0 | 1 |
| 379 | 3 | 0 |
| 388 | 3 | 1 |
| 389 | 2 | 2 |
| 402 | 3 | 2 |

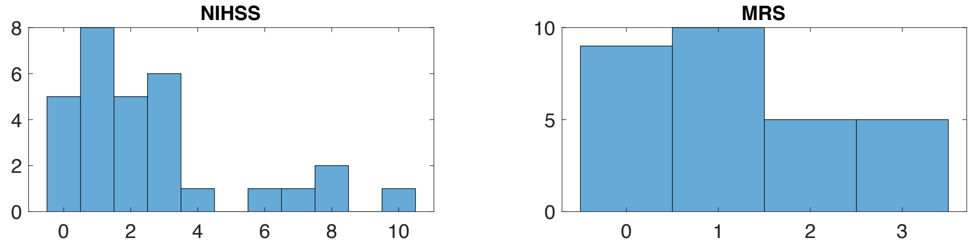


*Supplementary Figure 2. Distribution of the clinical scores for the National Institutes of Health Stroke Scale (NIHSS) and the modified Rankin scale (MRS)*

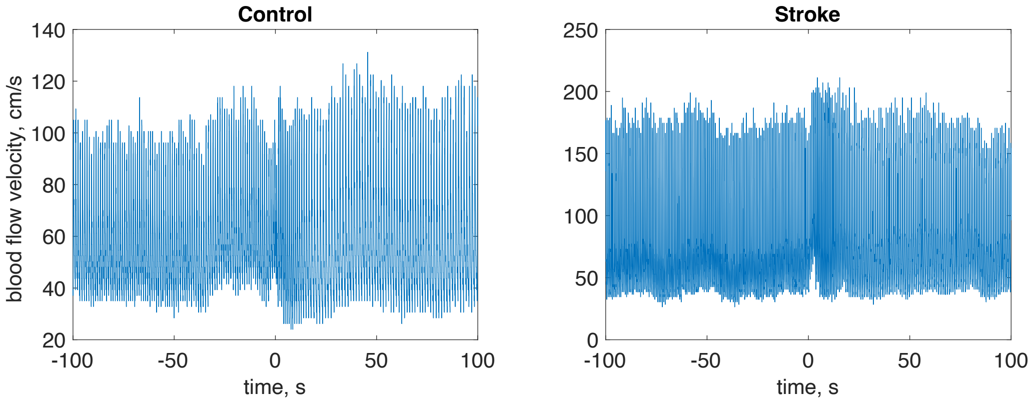

*Supplementary Figure 3. Samples of raw blood flow velocity measurements during the transition from sitting to standing (t=0) in a control and in a stroke patient.*

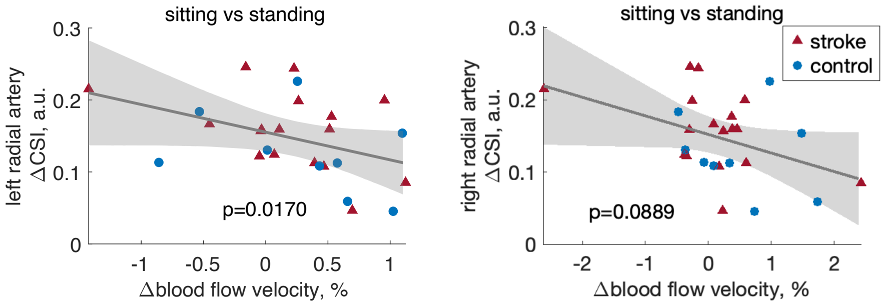


*Supplementary Figure 4. Control on the relationship between changes in blood flow velocity and cardiac sympathetic index (CSI), in the left and right radial arteries.*
